# Enrichment of helminth mitochondrial genomes from faecal samples using hybridisation capture

**DOI:** 10.1101/2025.02.24.639427

**Authors:** Marina Papaiakovou, Andrea Waeschenbach, Roy M. Anderson, Piet Cools, Zeleke Mekonnen, D. Timothy J. Littlewood, Cinzia Cantacessi, Stephen R. Doyle

## Abstract

New approaches are urgently needed to enrich rare or low-abundant DNA in complex samples. Soil-transmitted helminths (STHs) inhabit heterogeneous environments, including the gastrointestinal tract of their host as adults and are excreted as eggs and larvae in faeces, complicating our understanding of their biology and the use of genetic tools for species monitoring and population tracking. We have developed a hybridisation capture approach to enrich mitochondrial genome sequences of two STH species, the roundworm *Ascaris lumbricoides* and whipworm *Trichuris trichiura,* from extracted DNA from faecal material and worm specimens. Employing ∼1000 targeted probes, we achieved >6,000 and >12,000 fold enrichment for *A. lumbricoides* and *T. trichiura,* respectively, relative to direct whole genome shotgun (WGS) sequencing. Sequencing coverage was highly concordant with probe targets and correlated with the number of eggs per gram (EPG) of parasites present, from which DNA from as few as 336 EPG for *Ascaris* and 48 EPG for *Trichuris* were efficiently captured and sufficient to provide effective mitochondrial genome data. Finally, allele frequencies were highly concordant between WGS and hybridisation capture, suggesting little genetic information is lost with additional sample processing required for enrichment. Our hybridisation capture design and approach enables sensitive and scalable STH mitochondrial genome sampling from faecal DNA extracts and paves the way for broader hybridisation capture-based genome-wide applications and molecular epidemiology studies of STHs.

## 1. Introduction

Studying the genetic makeup of many organisms requires the isolation of the target DNA from a heterogeneous mixture of other contaminating nucleic acids. This is particularly true for the study of gastrointestinal (GI) parasitic worms, which exist and are in close interaction with the host, its microbiome, other parasitic species, and, potentially, the environment. Understanding these interactions is challenging, especially when DNA from the species of interest is significantly underrepresented in the mixture of nucleic acids that comprise the biological sample. Current detection of infections by GI parasitic worms relies on the morphological identification of microscopic eggs or larvae in stool samples (Zendejas-Heredia et al., 2021), a biological matrix predominantly composed of non-parasite material. More recently, molecular techniques such as quantitative PCR (qPCR) have increased the sensitivity of the detection of DNA from selected parasites in faecal extracts and have been applied to the detection of parasite infections based on the amplification of very short, species-specific DNA targets (Papaiakovou et al., 2021). However, current microscopy and qPCR-based diagnostic approaches provide little insight into the interactions among parasitic worms of the same species or into parasite-specific genetic traits, population dynamics, or responses to treatment. Furthermore, there is a lack of “information-rich” genetic tools to specifically target parasite DNA among complex samples.

There is an ongoing need to understand the genetics of parasitic worms and their interactions with their environment. One area where genetics may have value is to support the sustainable control soil-transmitted helminths (STHs). The World Health Organisation’s 2030 roadmap for Neglected Tropical Diseases (NTDs)(*Ending the Neglect to Attain the Sustainable Development Goals: A Road Map for Neglected Tropical Diseases 2021–2030*, 2021) aims to control parasite infections, including STHs, via large-scale deworming programs, coupled with the application of effective diagnostics to monitor interruption of transmission and eventually, elimination as a public health problem. Genetics may provide new insights into parasite traits relevant to their control, such as understanding transmission dynamics within and between populations (Durrant et al., 2020; Small et al., 2019), measuring population changes due to interventions (Hedtke et al., 2019), or monitoring genetic change under the selective pressures created by continual drug treatment (Doyle & Cotton, 2019). Despite an increasing number of studies on the genetics of STH based on the analysis of DNA from adult worms (Doyle et al., 2022; Easton et al., 2020; Landeryou et al., 2024; Liu et al., 2024), the ethical and logistical challenges of accessing adult specimens from humans represent a significant obstacle for large-scale cross-sectional or longitudinal studies. Meanwhile, recent studies have evaluated the application of whole genome sequencing (WGS) approaches on faecal samples for detecting and characterising the genetic diversity of helminths in stool samples (Papaiakovou et al., 2023, 2024). While mixed-species infections could be detected in some instances, genetic diversity analyses were restricted to highly fecund and abundant species (e.g. *Ascaris*), with limited or missing genetic data from poorly represented species. Moreover, direct DNA sequencing extracted from faecal samples yielded large numbers of bacterial DNA reads, significantly reducing the number of parasite reads available for downstream analyses. Therefore, new approaches are urgently needed to enrich low-abundant parasite DNA amongst complex and abundant off-target DNA.

Here, we evaluate hybridisation capture for enriching mitochondrial genomes of helminths from faecal DNA extracts. In this technique, single-stranded ‘probe’ sequences are hybridised to target regions within a DNA library. Unbound sequences are discarded via washing, resulting in an enriched DNA library of targeted sequences for sequencing. This approach has been used successfully in several parasitological studies, for example, to amplify the malaria parasite *Plasmodium falciparum* from human blood spots (Melnikov et al., 2011), to isolate coding regions by exome capture of the blood fluke *Schistosoma mansoni* to investigate the emergence of drug resistance (Chevalier et al., 2014; Le Clec’h et al., 2021) and species hybrids (Platt et al., 2019), and to sample *Leishmania* apicomplexan parasites from clinical samples (Domagalska et al., 2019). Using faecal DNA extracts from samples positive for two STH species, the roundworm *Ascaris lumbricoides* and whipworm *Trichuris trichiura,* we compare hybridisation capture versus WGS, focusing on target specificity, target enrichment, the sensitivity of parasite detection, and the ability to detect genetic variants.

## 2. Materials and Methods

### 2.1. Target selection and probe design

The primary objective of the hybridisation capture was to target and enrich the DNA of the mitochondrial genomes of *A. lumbricoides* and *T. trichiura*. Nuclear diagnostic repeats and internal transcribed spacers 1 and 2, were also targeted, albeit protocols associated with the latter are described in the Materials and Methods and associated Supplementary data but are not described in the main text. A total target size of 30,395 bp was provided to Daicel Arbor BioSciences (Ann Arbor, United States) for probe design. Targets were soft masked for simple and low-complexity repeats, after which 80-base probes in a 4× tiling array (i.e. one probe every ∼20 bp) were designed, resulting in 1,425 putative bait sequences.

### 2.2. Species-specificity of probe design

The specificity of the probes was tested using BLAST searches against the nuclear and mitochondrial genomes of other helminth species commonly present in faeces, including *Ancylostoma ceylanicum* (nuclear genome: PRJNA72583; mitochondrial genome: NC_035142.1), *Ancylostoma duodenale* (nuclear genome: PRJNA72581; mitochondrial genome: NC_003415.1), *Echinococcus multilocularis* (nuclear genome: PRJEB122; mitochondrial genome: NC_000928.2), *Enterobius vermicularis* (nuclear genome: PRJEB503; mitochondrial genome: NC_056632.1), *Necator americanus* (nuclear genome: PRJNA72135; mitochondrial genome: NC_003416.2), *S. mansoni* (nuclear genome: PRJEA36577; mitochondrial genome: NC_002545.1), *Strongyloides stercoralis* (nuclear genome: PRJEB528; mitochondrial genome: NC_028624.1), *Taenia solium* (nuclear genome: PRJNA170813; mitochondrial genome: MW718881.1). After filtering probes to minimise off-target binding, 1,076 of the original 1,425 baits were retained.

### 2.3. Probe coverage of target sequences

To determine the probe coverage on the two mitochondrial genome target sequences, the probe sequences were mapped using *minimap2* v.2.17-r941 to the mitochondrial and whole genome assemblies of *T. trichiura* (Doyle et al., 2022) and *A. lumbricoides (Easton et al., 2020)* (Supplementary Table 2). Unmapped sequences were removed with *samtools view* v.1.6 (Danecek et al., 2021), and *bedtools bamtobed* v2.30.0 (Quinlan, 2014) was used to convert the BAMfile to a BED file that included the mapped coordinates. The coordinates were sorted and merged, and the final BED file was used to visualise them (Supplementary Table 1). *Bedtools subtract* v2.30.0 was used to obtain the between probe/gap coordinates.

### 2.4. Sample selection and DNA extraction

A total of 24 biological samples were included in the study, including 23 faecal DNA extracts and one *A. lumbricoides* worm DNA extract (Supplementary Table S3). The collection and testing of parasite DNA from faecal samples was approved by Imperial College London, UK (Ref: 17IC4249 and 17IC4249 NoA1). The use of the DNA extract from the worm sample was approved by the Ethical Review Committee, Faculty of Medicine and Health Sciences / University Hospital, Ghent University, Belgium (Ref: B670201627755 and PA2014/003), and Jimma University, Ethiopia (Ref: RPGC/547/2016).

Of the faecal samples, 8 and 14 were positive for *Ascaris* and *Trichuris* by Kato-Katz (KK), respectively. DNA was extracted using the FastDNA SPIN kit for soil from MP Biomedicals (Santa Ana, CA) and a high-speed homogeniser with modifications, including an extraction control as described (Dunn et al., 2020; Papaiakovou et al., 2024). DNA from the single worm sample was extracted using the Isolate II Genomic DNA extraction kit (Bioline, Meridian Bioscience) (Papaiakovou et al., 2024). Thirteen and 18 of the faecal DNA extracts were positive for *Ascaris* and *Trichuris*, respectively, by qPCR (Supplementary Table 3).

### 2.5 Library preparation, hybridisation capture, and sequencing

DNA extracts were shipped to Daicel Arbor BioSciences, USA, for library preparation, hybridisation capture, and sequencing. After quantification and quality control checks, dual-indexed Illumina-compatible libraries targeting an average insert size of ∼300 bp were generated. For hybridisation capture, 250 ng of up to eight libraries per reaction were pooled for a total of three pools. Each hybridisation capture pool was dried down to 7 µL by vacuum centrifugation. Hybridisation captures were performed following the myBaits v5.02 protocol using a myBaits custom design (myBaits design ID: D10573KRILL) with an overnight hybridisation and wash at 65°C. Post-capture, half of the volume of the reactions was amplified for ten PCR cycles and subsequently quantified using a spectrofluorimetric assay. For hybridisation captures that did not yield sufficient material for sequencing, the remaining volume was amplified for 14 cycles.

Hybridisation captures were visualised using the Agilent TapeStation 4200 platform with a High Sensitivity D1000 tape. The captures were pooled in approximately equimolar ratios. Samples were sequenced using the Illumina NovaSeq 6000 platform on a partial S4 PE150 lane to achieve approximately two million read pairs per library. Each Illumina library was also sequenced directly (without hybridisation capture processing) to generate whole genome shotgun (WGS) data for comparative analyses. Libraries were pooled in equimolar ratios and sequenced on a partial NovaSeq X Plus instrument lane with 10B PE150 chemistry, targeting approximately ten Gb of data per sample.

### 2.6. Mapping of whole-genome and capture sequencing data

Demultiplexed raw sequencing reads from each dataset (hybridisation capture and WGS) were trimmed using *Trim-galore* v0.6.2 (Martin, 2011). To assess on-target enrichment and off-target contamination, trimmed sequencing reads from each dataset were each mapped to the metagenomic reference sequences (described above) using Burrow-Wheeler Aligner (BWA) Maximum Exact Match (MEM) v.0.7.17-r1188 (Li, 2013).

Unmapped reads were filtered using *samtools* v.1.6 *view* (Danecek et al., 2021), retaining reads >80 bp in length. Soft-clipped alignments were filtered by removing any reads with ‘S’ (indicating a soft-clipped alignment) or ‘H’ (indicating a hard-clipped alignment) on the CIGAR column of the BAM files. Lastly, duplicated reads were marked and removed from the BAM files using *Picard MarkDuplicates* v.picard-2.18.29-0 (Toolkit, 2019) and the number of reads was calculated using *samtools* v.1.6 *idxstats*. The percentage of total mapped reads to the target sequences was calculated using *samtools* v.1.6 *flagstat* (Danecek et al., 2021) and was visualised in R v.4.2.2.

The number of mapped reads per sample per species was recovered using *bedtools* v.2.30 *multicov (Quinlan, 2014)* on filtered BAM files. To compare samples, read count normalisation was performed as follows:

*normalised reads = reads mapped / total reads / mitochondrial genome size (Mb)*

### 2.7. Target enrichment using hybridisation capture versus WGS

To assess the efficacy of target enrichment compared to the WGS data, we calculated the absolute (total normalised reads) and relative (ratio of capture:WGS reads) fold enrichment. This was determined by comparing the two datasets’ normalised read counts per sample for each species. A higher fold enrichment value indicates a greater normalised read count mapping to the targets in the hybridisation capture than the corresponding sample in the WGS datasets.

### 2.8. Per sample depth of coverage in probe-binding regions and gaps

*Samtools depth* v.1.6 was used to calculate the per-base coverage across all samples (with maximum allowing depth -d 1000000). Per base read depth was normalised by the median read depth across the entire length of the mitochondrial genome, per sample per species. In *Ascaris*-positive samples only, the depth of coverage in the regions not covered by probes (= gaps) was calculated similarly using *samtools depth* v.1.6, and per base depth was normalised by the median depth per sample ID, as described above. Then, each normalised gap depth was further normalised by the maximum gap median depth (resulting in a maximum depth of 1) for direct comparison between different samples. No gap coverage was calculated for *T. trichiura*, as the probes covered the entire length of the mitochondrial genome except for two gaps of 20 bp each.

To assess genome coverage, we analysed the proportion of the mitochondrial genome covered at >1X, >10X, and >100X for hybridisation capture and WGS datasets. This was achieved by calculating the proportion of bases in the mitogenome with a read depth exceeding each threshold using *samtools depth* v.1.6.

### 2.9. Relationship between helminth egg burden and sequencing depth in faecal samples

To investigate the relationship between parasite load and sequencing depth, eggs per gram (EPG) in all faecal samples (n = 23) were compared with the mean sequencing depth. The relationship between EPG and mean sequencing depth was analysed using Spearman’s rank correlation (p.accuracy = 0.001, r.accuracy = 0.01), a non-parametric method well-suited for capturing relationships between two variables that may not be linear. This approach was aimed to determine the minimum number of eggs required for effective hybridisation capture or WGS, which we defined in microscopy-positive samples were > 50% of the mtDNA was covered with sequencing reads. This threshold highlights a critical benchmark for sequencing success in samples with varying parasite loads.

### 2.10. Variant calling in mitochondrial genomes

Variant calling was performed using *bcftools mpileup* (v.1.16) (Danecek et al., 2021) to generate a Variant Calling Format (VCF) file containing only variant sites (used for pooled data, downstream, filtered for both min and max number of alleles = 2). To account for variants that were present in single, unpaired reads, the following code was applied: *bcftools mpileup -A --annotate FORMAT/AD -Ov -f REF.fasta -d 100000 -b bamlist_for_variant_call | bcftools call --ploidy 1 -mv --skip-variants indels -o CAPTURE_DATA_ALL_SAMPLES_bcftools_with_orphan_reads.vcf*. Variants were further filtered to ensure (i) a maximum of two alleles, (ii) unique variants, and (iii) only SNPs (no insertions/deletions) were retained. The VCF files were converted into data frames in R using vcf2Rtidy (*vcfR* v.1.15.0). Allele frequencies were calculated as ALT_depth/SUM_depth for each SNP. SNPs detected by both hybridisation capture and WGS data were identified, and their correlation was assessed using Pearson’s method (confidence level = 0.99).

### 2.11. Data availability

The final set of probe-target coordinates and sequences are provided in Supplementary Table 1. Reference genomes used for read mapping and comparative genomics are described in Supplementary Table 2. Study and sample accessions for the sequencing data generated and analysed in this study are described in Supplementary Table 3.

### 2.12. Code availability

The code used in the study is available via GitHub at: https://github.com/MarinaSci/STH_HYBRIDISATION_CAPTURE.git.

## 3. Results

### 3.1. Hybridisation capture enriches *Ascaris* and *Trichuris* DNA

To test our approach for enriching helminth mitochondrial genomes from faecal DNA extracts, we sequenced hybridisation capture and WGS libraries from 23 faecal DNA extracts and one whole worm DNA sample (expelled from an infected person).

Sequencing and read processing yielded 2.6 billion (range: 61.4 - 189.1 million per sample) and 108.4 million (range: 60,020 - 30.7 million per sample) trimmed paired-end reads for WGS and hybridisation capture, respectively (Supplementary Table 4).

We first compared the effectiveness of hybridisation capture for enriching target sequences relative to WGS by assessing the proportion of reads successfully mapped to target regions (on-target) versus reads not mapped to any of the targets (off-target). Hybridisation capture resulted in a mean percentage of 11.94% on-target reads from all faecal samples (Figure 1A; range = 0.52 - 48.99%, 16.41% standard deviation (SD)). In comparison, the mean percentage of WGS reads mapped was 0.01% (Figure 1A; range = 0-0%; SD = 0.00%). For reads obtained from adult worm DNA, 9.94% and 99.10% of WGS and hybridisation capture reads, respectively, were successfully mapped to target regions. Mapping to non-target regions by the remaining reads in both datasets was attributed to non-specific binding of the probes to off-target sequences (e.g. bacterial), as revealed by BLAST analysis.

**FIGURE 1.**
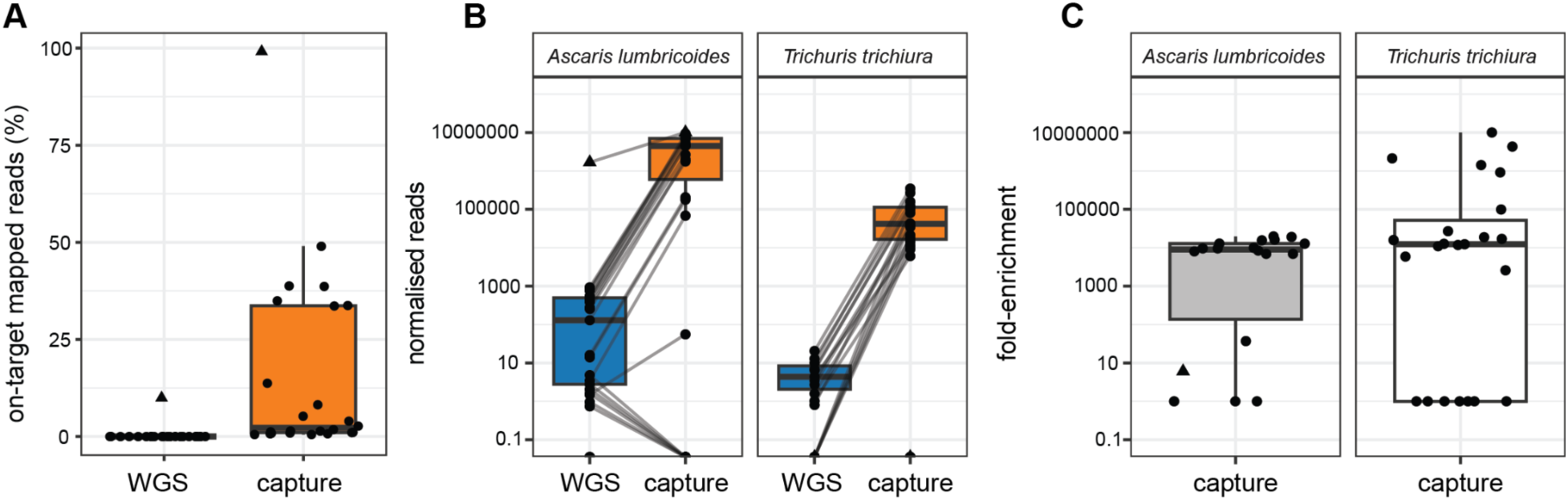
Hybridisation capture efficiency of targeting helminth mitochondrial genomes in faecal DNA extracts. (A) Boxplots indicating on-target mapped reads (%) for whole genome shotgun (WGS) (blue) and hybridisation capture (orange) of DNA from faecal and worm DNA extracts for both *Ascaris lumbricoides* and *Trichuris trichiura*. (B) Boxplots indicating enrichment based on normalised (defined as reads mapped per million trimmed reads per Mb) for WGS (blue) and capture (orange) in faecal and worm DNA extracts for *A. lumbricoides* and *T. trichiura*. Grey lines connect corresponding DNA extracts across WGS and hybridisation capture. (C) Boxplots indicating fold enrichment (calculated as fold enrichment = normalised reads from hybridisation capture / normalised reads from WGS) from hybridisation capture data for *A. lumbricoides* and *T. trichiura*. In each boxplot, the central box represents the interquartile range (IQR; from Q1 and Q3), with the median displayed as a line through the centre of the box. The whiskers extend 1.5 times the IQR from Q1 and Q3, respectively. In all plots, DNA extracts from adult *Ascaris* are represented by a triangle, whilst circles represent faecal DNA extracts.

The degree of enrichment between the hybridisation capture and WGS datasets was assessed by determining the normalised read count mapped to each mitochondrial genome target. Normalised reads were defined as the number of reads mapped per million total trimmed reads per megabase (Mb) of the mitochondrial genome per sample. For the faecal WGS data, the median normalised reads were 14 and 0 for *A. lumbricoides* and *T. trichiura*, respectively, whereas the hybridisation capture dataset yielded median normalised read counts of 178,499 and 17,543, respectively, thus indicating a significant enrichment in the hybridisation capture dataset compared to WGS (Figure 1B; Supplementary Table 4). For adult *Ascaris*, WGS and hybridisation capture data yielded a median of 1,676,426 and 10,109,604 normalised reads, respectively. Whilst six of the 23 faecal DNA extracts did not show enrichment in hybridisation capture relative to WGS, each of these extracts was negative for *A. lumbricoides* by both KK and species-specific qPCR, and yielded >10 normalised reads by WGS, indicating very low parasite burdens. BLAST analysis of a subset of these reads returned positive hits for *A. lumbricoides*; nevertheless, the reads mapped to regions not covered by probe design or areas of probe coverage intercalated with gaps.

Finally, we determined the relative fold-enrichment of the hybridisation capture compared with WGS, calculated as the ratio of normalised reads yielded by hybridisation capture divided by that yielded by WGS (Figure 1C; Supplementary Table 5). This ratio highlights the proportional improvement in target enrichment achieved through hybridisation capture, making it easier to compare the relative efficiency across different samples. The median enrichment for *A. lumbricoides* and *T. trichiura* targets in faecal DNA extracts was 6,931-fold and 12,347-fold, respectively. In contrast, hybridisation capture performed on DNA from adult *Ascaris* yielded a six-fold enrichment only, likely due to the greater efficiency of WGS on purified worm DNA.

### 3.2. Hybridisation capture yields consistent mitochondrial genome sequencing coverage

Next, we explored the relationship between hybridisation probe binding sites and sequencing coverage by comparing mitochondrial protein- and ribosome-coding gene features (Figures 2A, 2E), contiguous regions of probe coverage (Figures 2B, 2F), as well as gaps in probe coverage (Figures 2C, 2G), to the normalised depth per faecal DNA extract across the whole mitochondrial genome sequence (Figures 2D, 2H) of both *A. lumbricoides* and *T. trichiura,* respectively. The equivalent comparison for the single *A. lumbricoides* worm is shown in Figure S1.

**FIGURE 2.**
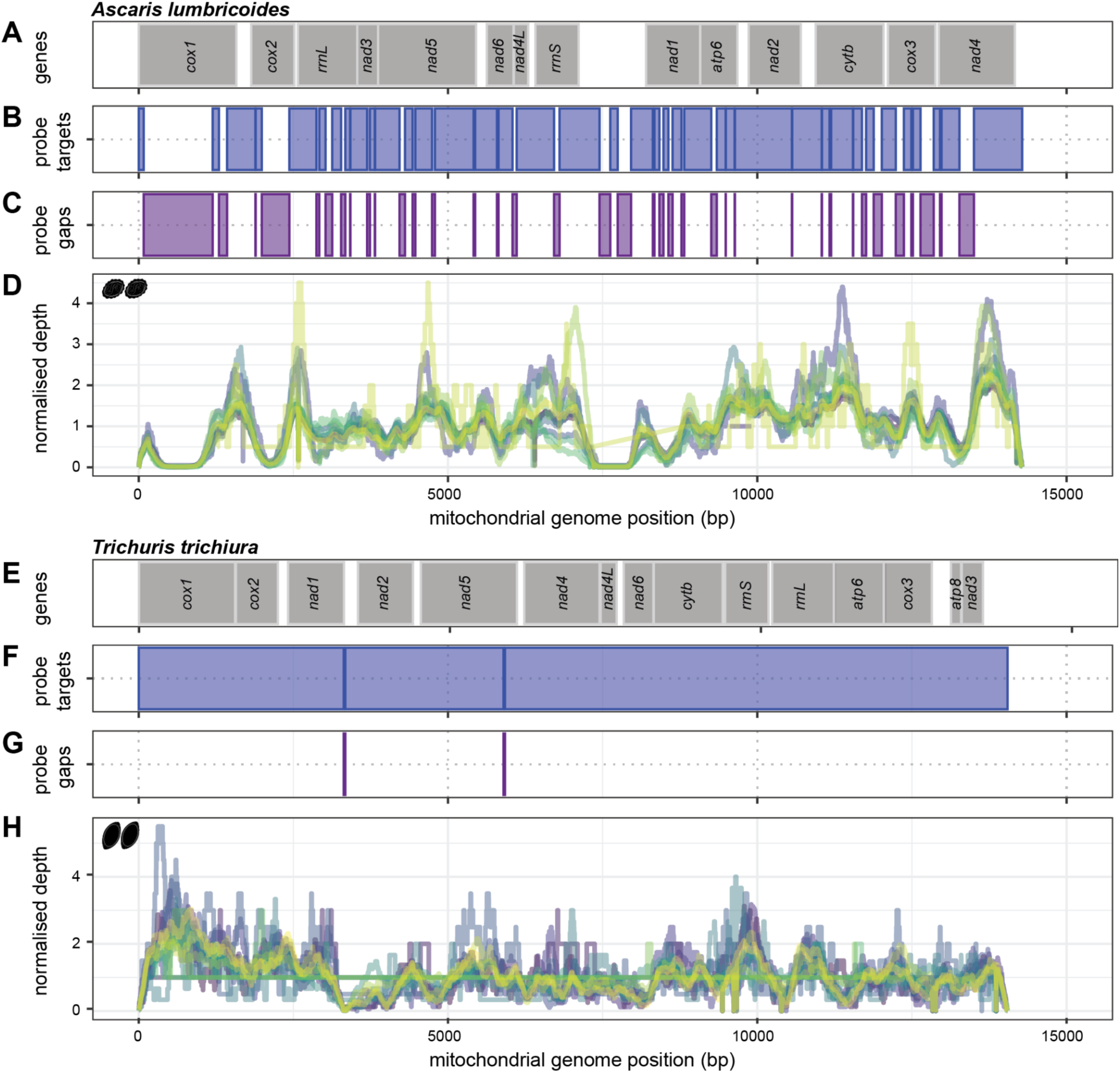
Relationship between hybridisation probe binding sites and sequencing coverage across mitochondrial genomes. We compared protein- and ribosome-coding mitochondrial gene coordinates (A and E), hybridisation probe coverage (B and F), gaps in probe coverage (C and G) against normalised read depth for *Ascaris lumbricoides* (n = 14) (E) and *Trichuris trichiura* (n = 16) (H), respectively. Line colours in the normalised depth plots represent individual DNA extracts. The normalised read depth was calculated by dividing the per base read depth by the median read depth of the genome; a null expectation of even coverage would result in a normalised read depth of one.

A consistent relationship between probe and sequencing coverage was observed, particularly highlighted by regions of no probe coverage and a decrease in sequencing depth. This was particularly evident in the *A. lumbricoides* data, which were characterised by large gaps in probe coverage across *cox*1 and *cox*2 and the control region, as indicated by a large space between genes toward the middle of the mitochondrial genome (Figure 2C). For *T. trichiura*, sequencing coverage was consistent across all DNA extracts due to the greater coverage of probes. Higher variation in coverage was observed in the hybridisation capture data versus the WGS, including in data from adult *Ascaris* (Figure S1), likely as a consequence of the additional processing steps required for hybridisation capture library preparation and inherent variation in probe binding, density, and GC content between regions.

Nevertheless, the absence of probes targeting any given region did not result in a complete lack of sequencing depth but rather in decreased coverage in gap regions. To assess the relationship between gap length and coverage, we calculated a normalised per base coverage of each gap (bases mapped / gap length / median genome-wide coverage) (Figure S2). This analysis revealed that gaps <25 bp were typically fully covered at maximum depth and decreased in coverage as gaps increased in length with 50% of maximum depth up to a gap of approximately 250 bp (Figure S2). These findings highlight that additional data are obtained even in non-targeted regions adjacent to probe targets.

### 3.3. Hybridisation capture results in concentration-dependent target coverage

A motivating factor for developing an enrichment approach for STH from faecal samples is to enable genomic analyses of sequence variation in targeted sequences. Depending on the downstream analysis, minimum sequencing coverage thresholds are typically applied. We explored the effective coverage, defined as a proportion of the length of the genome with mapped sequencing reads, at depth thresholds of >1X, 10X, and 100X for *A. lumbricoides* (Figures 3A, 3B) and *T. trichiura* (Figures 3C, 3D). Hereafter, “effective coverage” is defined as achieving successful read mapping across >50% of the whole mitochondrial genome.

**FIGURE 3.**
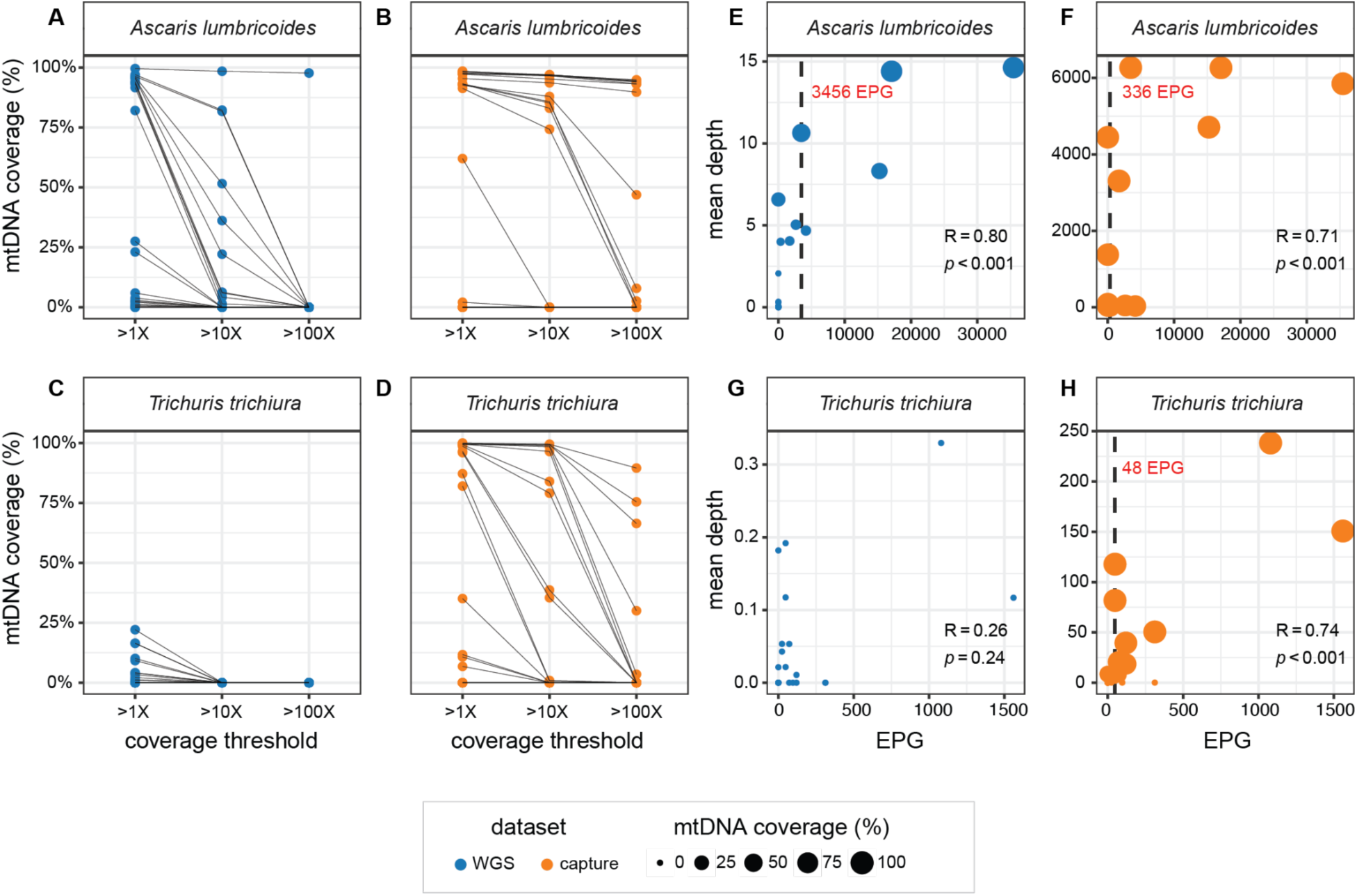
Comparison of effective mitochondrial coverage at sequencing depth thresholds of >1x, >10 and >100x and relationship to parasite eggs per gram (EPG) in faecal samples. (A - D) Effective target coverage, defined as the percentage of the mitochondrial genome covered by sequencing reads generated by WGS (blue) and hybridisation capture (orange) at sequencing depth thresholds of >1X, >10X and >100X, respectively, for (A and B) *Ascaris lumbricoides* positive samples and (C and D) *Trichuris trichiura* positive samples. In each plot, circles represent individual samples, and grey lines connect the same sample across different depth thresholds. (E - H) The relationship between sequencing depth and parasite burden is represented by the number of eggs per gram (EPG) in WGS (blue) and hybridisation capture (orange) data for *A. lumbricoides* positive samples (E and F) and *T. trichiura* positive samples (G and H). In each plot, the diameter of each circle depicts the effective target coverage of each sample, as defined above. Spearman’s correlation coefficient and associated p-value comparing EPG and mitochondrial genome depth are shown. The vertical dashed black lines depict the minimum EPG to achieve >50% of mitochondrial genome coverage. No minimum EPG could be calculated for the WGS data for *T. trichiura*.

Of the 23 faecal DNA extracts for *A. lumbricoides,* 14 and seven were positive by qPCR and KK, respectively (Supplementary Table 3). For *A. lumbricoides*, effective coverage was achieved in 11, four, and one sample by WGS at >1X, >10X, >100X coverage thresholds, respectively (Figure 3A), and 14, 13, and eight, respectively, by hybridisation capture (Figure 3B). For *T. trichiura*, 18 and 14 samples were positive by qPCR and KK, respectively (Supplementary Table 3). No DNA extracts yielded effective coverage by WGS (Figure 3C), whilst 12, eight, and three samples achieved effective coverage by hybridisation capture (Figure 3D). For both species, hybridisation capture resulted in more samples with greater than 50% effective coverage at a greater range of sequencing depths than WGS.

For STHs, parasite EPG is frequently applied as a proxy of infection intensity and the quantity of parasite DNA in a given sample. We determined the relationship between EPG and sequencing depth and the minimum EPG required to achieve effective coverage of the mitochondrial genomes of both *A. lumbricoides* and *T. trichiura*. With one exception (WGS of *T. trichiura*), a significant positive relationship was observed between EPG and sequencing depth (R = 0.80, p < 0.001 and R = 0.71, p < 0.001 for *Ascaris* for WGS and hybridisation capture, respectively; R = 0.74, p < 0.001 for *Trichuris* hybridisation capture). For *Ascaris,* the minimum EPG required to achieve effective coverage by WGS was 3,456 (Figure 3E) and 336 by hybridisation capture (Figure 3F). For *Trichuris,* no effective coverage could be achieved by WGS (Figure 3G), whereas the minimum EPG required to achieve effective coverage by hybridisation capture was 48 (Figure 3H).

### 3.4. Variant allele frequencies are highly correlated between WGS and hybridisation capture

Concordant genetic data between WGS and hybridisation capture approaches are crucial for ensuring methodological robustness and drawing meaningful conclusions from biological signals. We next compared the number of genetic variants detected by each or both WGS and hybridisation capture. Variant calling revealed 510 and 1,406 SNPs for *Ascaris* and *Trichuris* by hybridisation capture and 490 and 481, respectively, by WGS (Supplementary Table 4). For *Ascaris*, 90.92% of genetic variants were detected by both approaches, whilst 6.5% were uniquely detected by hybrid capture and 2.68% by WGS (Figure 4A). In contrast, for *Trichuris*, only 29.20% of SNPs were detected by both approaches, with most SNPs (67.10%) solely detected by hybridisation capture (Figure 4A), consistent with the significantly higher enrichment of *Trichuris* DNA by hybridisation capture compared to WGS. To validate variants, a depth filter of >10 reads per SNP was set, which led to 499 SNPs for *Ascaris* and 1,406 for *Trichuris* by hybridisation capture and 484 (*Ascaris*) and 0 (*Trichuris*) by WGS (Figure 4B). Of note, all *Trichuris* SNPs detected by WGS were low-depth variants (Supplementary Table 7). Seventeen of the 19 SNPs uniquely detected by WGS data were in regions covered by the probes, whilst ten (out of 17) were in the control region of the mitochondrial genome.

**FIGURE 4.**
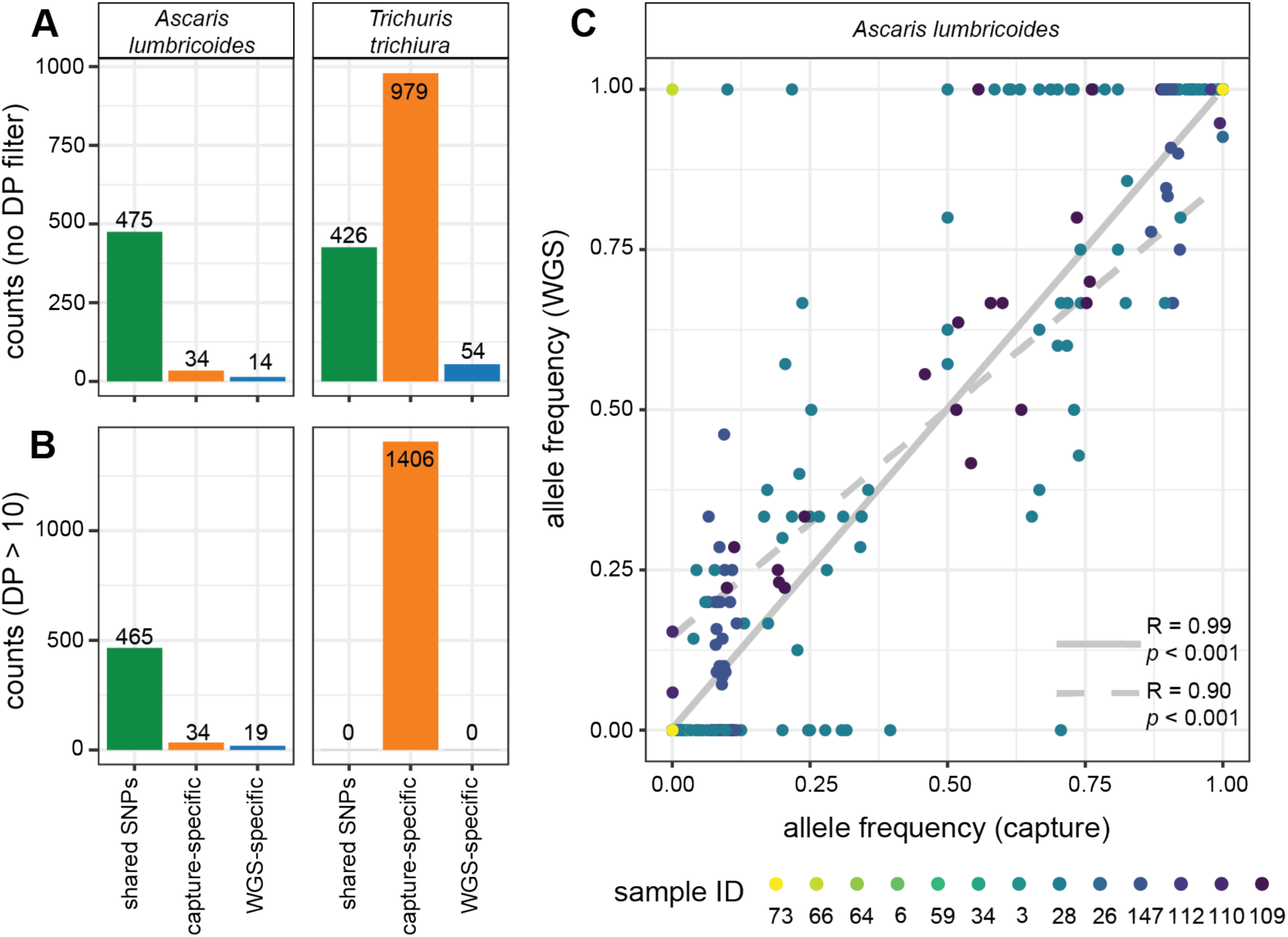
Comparison of genetic variation captured by WGS versus hybridisation capture. (A) Barplots show shared (green), capture-specific (orange) and WGS-specific (blue) SNPs for *Ascaris lumbricoides* and *Trichuris trichiura* before filtering for sequencing depth (DP). (B) Barplots show shared (green), capture-specific (orange) and WGS-specific (blue) SNPs for *A. lumbricoides* and *T. trichiura* following the selection of SNPs with DP > 10. (C) Comparison of allele frequency between SNPs identified by both hybridisation capture and WGS. Two Pearson’s correlation coefficients are shown, comparing (i) SNPs detected by both WGS and hybridisation capture (n = 460 SNPs, n = 13 individuals; solid grey line) and (ii) SNPs detected by both WGS and hybridisation capture, following filtering of allele frequencies of zero and one (n = 55 SNPs, n = 5 faecal extracts; dashed line).

Finally, to compare genetic concordance between WGS and hybridisation capture in *Ascaris*, we compared the allele frequencies of shared SNPs with sequencing depth (DP) > 10 (Figure 4B and 4C). This analysis was only performed on *Ascaris* data, given that no SNPs were shared between approaches for *Trichuris* (Figure 4B). A strong positive relationship in allele frequencies between variants detected by both hybridisation capture and WGS was observed (R = 0.99, p < 0.001; n = 460 SNPs, n = 13 individuals), mostly attributable to a large proportion of shared, fixed allele frequencies; following removal of variants with allele frequencies of 0 or 1 from both datasets, the strong positive correlation was retained (R = 0.90, p < 0.001; n = 55 SNPs, n = 5 individuals). These data suggest hybridisation capture, even with stringent filtering, can effectively capture the genetic variation present.

## 4. Discussion

Hybridisation capture of targeted DNA sequences together with high-throughput sequencing have become key tools for studying the genetics of low-abundant organisms in complex samples, enabling new insights in diverse research areas such as evolution and paleobiology (Gaudin & Desnues, 2018), drug resistance, and diagnostics (Melnikov et al., 2011), surveillance and epidemiology (Quek & Ng, 2024). To the best of our knowledge, hybridisation capture has not yet been applied to soil-transmitted helminths (Quek & Ng, 2024). Stool is a biological matrix frequently used to diagnose and characterise parasitic and other infections, as it can be collected non-invasively and analysed relatively inexpensively using basic laboratory equipment and expertise (Papaiakovou et al., 2022). However, the ability to diagnose infections can vary, particularly in cases characterised by low infection burdens and high levels of bacterial and host contamination. Here, we demonstrate that hybridisation capture leads to the enrichment of genomic targets of multiple STH from faecal DNA, providing the basis for a greater understanding of the genetic diversity and molecular epidemiology of parasitic worms directly from stool samples.

Genomic approaches are increasingly applied to the study of STH genetic diversity. Whole genome sequencing of adult parasites has been extensively applied to investigations of such genetic diversity or for epidemiological studies of ‘who infects whom’ (Doyle et al., 2022; Easton et al., 2020; Landeryou et al., 2024; Liu et al., 2024) however, worm specimens are not typically readily available or accessible, limiting their scope for large scale studies. Whole genome sequencing of faecal DNA extracts led to the recovery of genetic data from highly abundant helminth species, albeit efficiency was significantly reduced in cases of mixed-species or low-abundance infections (Papaiakovou et al., 2023, 2024). Although metabarcoding approaches are being developed (Chan et al., 2022; Kang et al., 2024; Miller et al., 2024; Papaiakovou et al., 2022; Venkatesan et al., 2025), they focus on very short target sequences that assist species identification but are less useful for characterising intraspecific genetic diversity. Our hybridisation capture approach applied directly to faecal DNA extracts significantly enriched the mitochondrial genomes of *Ascaris lumbricoides* and *Trichuris trichiura* by more than 6,000-fold and 12,000-fold, respectively, compared to WGS. For both species, hybridisation capture achieved coverage of the mitochondrial genomes in target capture regions despite increased variation in coverage relative to WGS. The strong correlation between the infection burdens (as measured by eggs per gram) and sequencing depth suggests that EPG can inform the efficacy and efficiency of the hybridisation capture, with hybridisation capture requiring ten times fewer eggs to achieve the same degree of mitochondrial genome completeness as WGS. As such, EPG may be used as a defining criterion for selecting samples suitable for hybridisation capture. As defined by the World Health Organisation, samples with <5,000 EPG for *A. lumbricoides* or <1,000 EPG for *T. trichiura* are classified as low/mild infection levels (Montresor et al., 2015); based on our findings, hybridisation capture represents an effective approach to monitor the genetics of these parasites in low-intensity settings, which may provide critical information for parasite control programs aiming to interrupt transmission.

Hybridisation capture offered substantial coverage of the mitochondrial genome, enabling the identification of most SNPs identified by WGS. In some cases (e.g. *Trichuris*), SNPs were only detected by hybridisation capture due to the insufficient coverage achieved by WGS. This finding suggests that hybridisation capture sequencing may be applied *in lieu* of WGS for studies of mitochondrial genome variation, providing equivalent information at a fraction of the sequencing coverage. This is of great importance in molecular epidemiological studies of how transmission persists in defined communities of people. Moreover, the demonstrated >6,000-12,000 fold target enrichment suggests that hybridisation capture is more cost-efficient than WGS per data unit despite a higher cost per sample. Some variation was missed by hybridisation capture data due to gaps in probe design. However, even taking into account the exclusion of some probes, gaps up to ∼ 25 bp are tolerated without compromising the overall coverage, whilst larger gaps between probe-covered regions may still receive some coverage up to 1,000 bp away from the targeted region, essentially providing additional capture data at no extra cost. More than half of the unaccounted *Ascaris* variants – detected by WGS but not by hybridisation capture – were located in the mitochondrial control region, a hypervariable non-coding region of the genome typically prone to mismapping and, as such, not usually included in downstream genetic analyses. Further design efforts to optimise probe positioning or sequencing library fragmentation size could enable more or less additional data as required.

There were limitations to our study. We observed substantial differences in probe coverage between species due to the selection of probes based on available genetic information on closely related species during the probe design. This step was likely redundant; keeping all probes and mapping the sequencing reads competitively to multiple reference sequences would allow non-specific but closely related sequences to align to their appropriate reference. In addition, whilst we determined the minimum EPG to achieve effective mitochondrial coverage, we acknowledge that such determination is based on a multi-copy target and will thus require validation based on other (single-copy) targets. Similarly, designing an efficient capture strategy to target multiple DNA sequences with different copy numbers, for example, multiple species that vary in fecundity, would need further validation. Costs associated with the hybridisation capture techniques depend on several factors, including probe pool size, number of samples pooled, and sequencing depth required. Thus, WGS sequencing applied to worm DNA extracts is comparatively more cost-efficient, as the relative fold enrichment was minimal compared to that obtained from faecal DNA extracts. The cost of hybridisation capture sequencing is also significantly higher than traditional diagnostic methods, and as such, it is not proposed to replace such tools but rather to acquire genetic information on circulating STHs.

In summary, hybridisation capture complements conventional diagnostic methods for characterising STH in complex biological samples such as faeces. The ability to efficiently characterise genetic variation for such parasites offers unprecedented insight into the diversity and evolution of parasitic organisms, thereby aiding our understanding of their infection patterns and dynamics. Such technologies may also assist in the detection of low-abundant parasites in other complex samples, such as water and soil. This could further aid management efforts to understand STH epidemiology and support efforts toward morbidity control and, ultimately, transmission elimination.

## Acknowledgements

This work was funded by an internal Natural History Museum (London) Science Investment Fund award. Samples provided by Ghent University, Belgium, were part of field trials supported by the Bill & Melinda Gates Foundation (USA; OPP1120972). MP is supported by a Harding Distinguished Postgraduate Scholarship held at the University of Cambridge, UK. SRD is supported by a UKRI Future Leaders Fellowship [MR/T020733/1] and the Wellcome Trust [206194]. The authors thank Nay Yee Wyine (London for Neglected Tropical Disease Research), Kay Thwe Han (Department of Medical Research, Ministry of Health and Sports, Nyapyitaw, Myanmar), Aye Moe Moe Lwin and Nay Soe Maung (University of Public Health, Myorma Kyaung Street, Yangon, Myanmar) for their help with the original epidemiological study and sample provision conducted in Myanmar. We appreciate Julia Dunn’s assistance in facilitating access to the samples from Myanmar and her direct contributions to sample processing. We thank Soe Thiha Lwin and Khine Khine Su (Defence Services Medical Academy, Pyay Road, Mingaladon, Yangon, Myanmar) and Myint Myint Sein (University of Public Health, Myorma Kyaung Street, Yangon, Myanmar) for their help with the fieldwork. We gratefully acknowledge Bruno Levecke for facilitating the availability of the sample from Ethiopia, Jennifer Klunk from Daicel Arbor Biosciences for technical support and advice, Matthew D. Clark and Oliver W. White for co-securing the funds with MP to deliver this project, Darren Chooneea for early discussions with MP on selective enrichment of targets, and the Helminth Genomics group at the Wellcome Sanger Institute for their support and feedback on the project.

## Data Accessibility and Benefit-Sharing

### Data accessibility statement

Raw sequence reads are deposited in the SRA (BioProject TBD).

### Benefit-sharing statement

The benefits of this research arise from our commitment to publicly sharing code, data, and results, as previously indicated.

## Author Contributions

MP conceived the study; MP and SRD designed the study; SRD supervised the study; RMA, PC, and ZM provided materials; MP prepared samples for sequencing, led the bioinformatics and analysed the results; MP and SRD interpreted the results, with input from contributions from AW, CC, and DTJL; MP drafted the original manuscript with input from SRD. All authors contributed to the revision and approval of the final paper.

