## Supplementary Information for "Enrichment of helminth mitochondrial genomes from faecal samples using hybridisation capture"

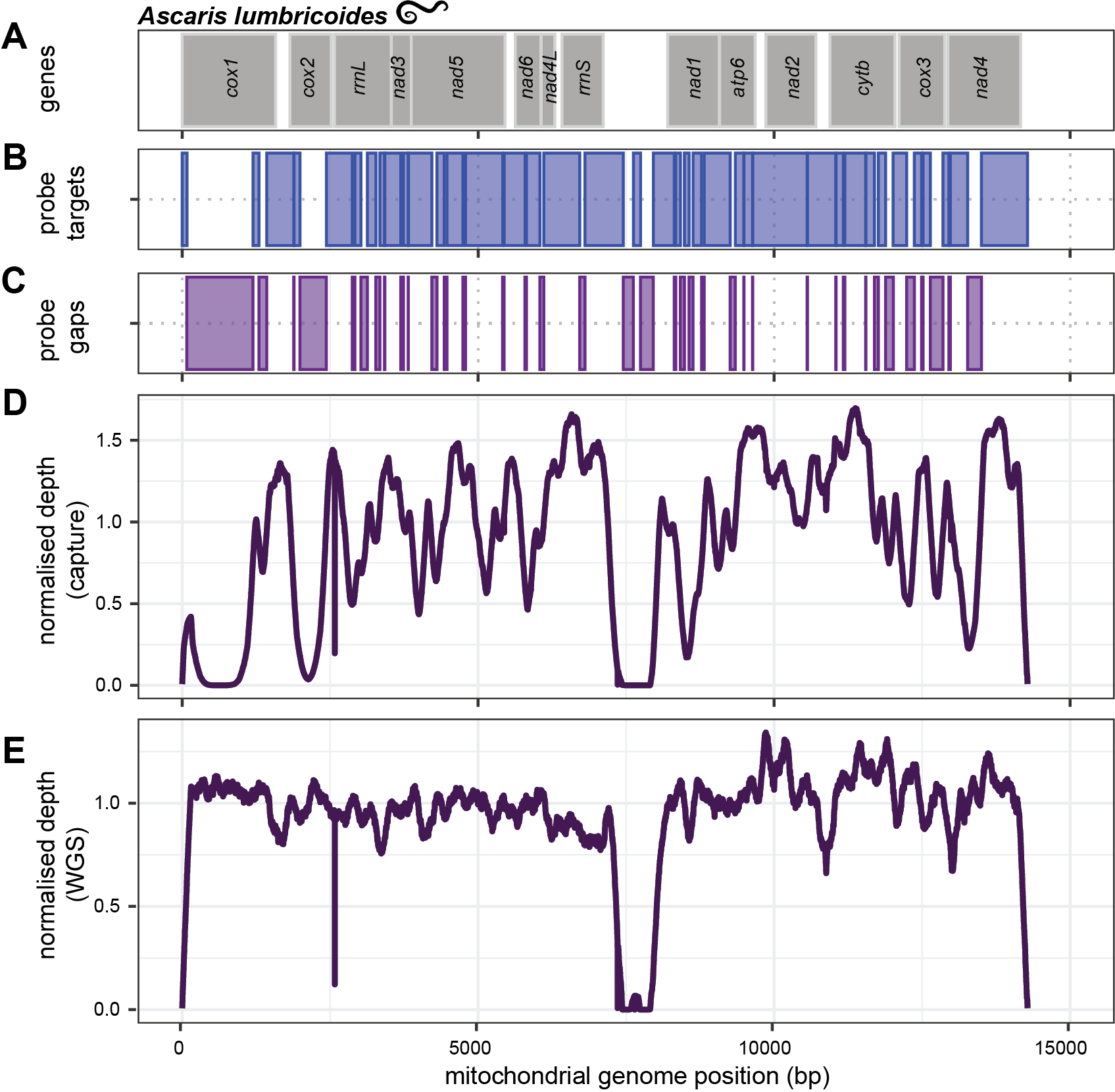
**Figure S1. Probe/bait, gap and mitochondrial genome coverage in *Ascaris lumbricoides* worm data from capture and WGS data.** (a) Boundaries of protein- and ribosome-coding mitochondrial genes for *A. lumbricoides* (worm)*.* (b) Coordinates of probe/bait targets for the mitochondrial genome of *A. lumbricoides* (worm)*.* (c) Coordinates of probe/bait gaps for the mitochondrial genome of *A. lumbricoides.* (d) Raw depth was normalised by the median depth, per sample, per species, across the mtDNA of *A. lumbricoides* from a single *Ascaris* worm obtained from capture*.* (e) Raw depth was normalised by the median depth, per sample, per species, across the mtDNA of *A. lumbricoides* from a single *Ascaris* worm obtained from WGS.

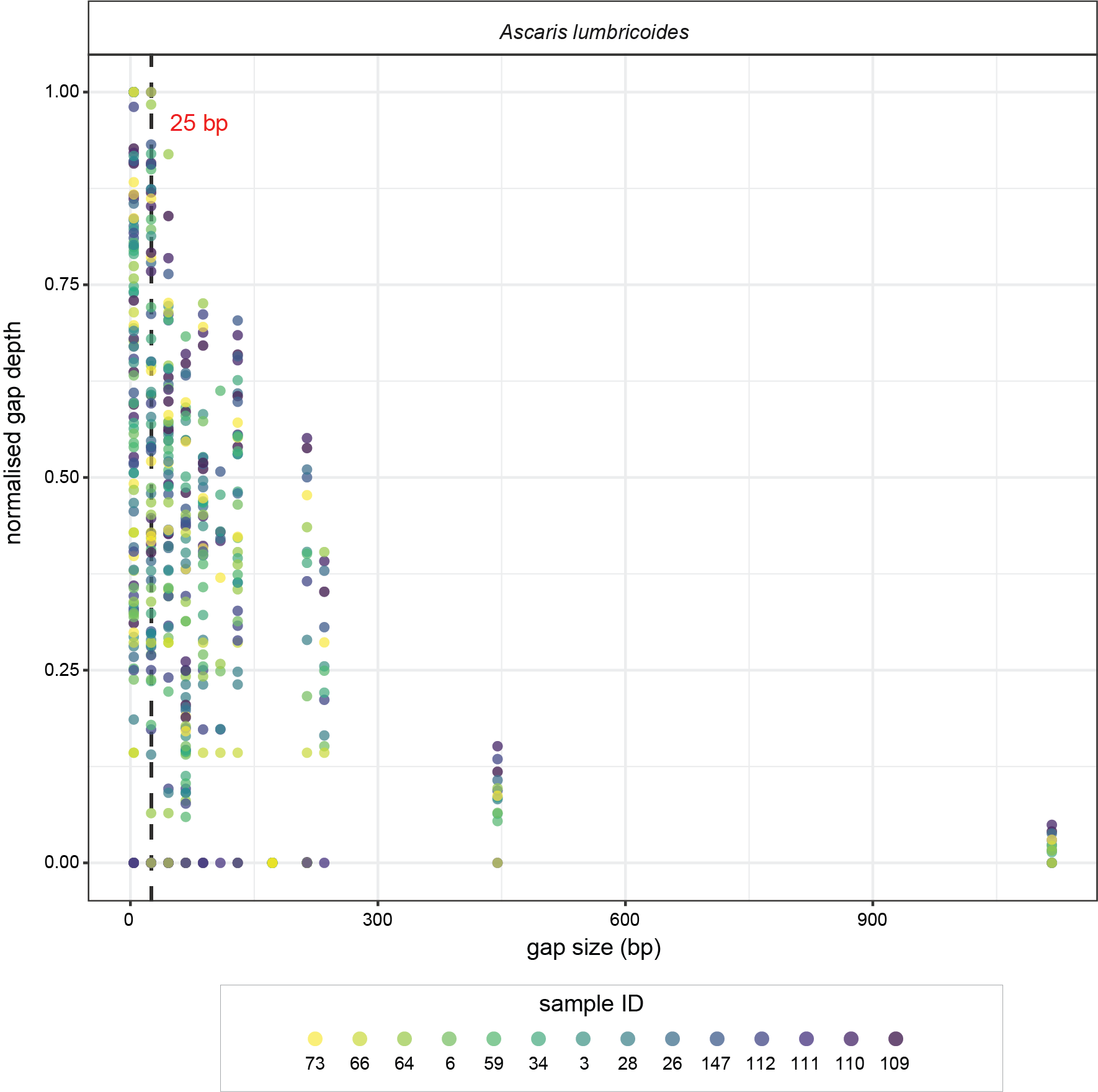

**Figure S2. Relationship between gap size (between probes/baits) and normalised gap depth.** Raw depth was calculated for the gap coordinates and normalised by the median depth per sample ID. Normalised depth was further normalised by dividing each value by the max value across all gaps for that sample ID. The black dashed line (25 bp) depicts the maximum gap value for the maximum normalised gap depth (=1).
